# Redundant contacts and force redistribution stabilize limbless vertical climbing

**DOI:** 10.64898/2026.07.06.736779

**Authors:** Calvin A. Riiska, Michelle Lee, Yonatan Nemenman, Gauge Thacker, Joseph R. Mendelson, Jennifer M. Rieser

## Abstract

Animals navigating complex vertical environments must secure stable footholds, a challenge for species without feet. While arboreal climbing has evolved repeatedly in snakes, the physical mechanisms they use to scale broad, nearly flat surfaces remain poorly understood. By measuring three-dimensional body kinematics and per-contact forces on a smooth vertical wall with protruding posts, we show that cornsnakes climb by dynamically balancing forces across a highly redundant network of 5 to 16 simultaneous contacts—far exceeding the three contacts minimally required for physical stability. Using a computational model and a robotic climber, we demonstrate that while simple body undulations and passive friction are mechanically sufficient to climb this terrain, snakes systematically deviate from this passive baseline. While downward climbing relies primarily on friction, ascending snakes actively generate positive mechanical work at their contacts to propel themselves. Furthermore, we found that whenever a snake engages a new contact, it triggers a stereotyped, system-wide redistribution of force that seamlessly integrates the new foothold without disrupting whole-body balance. These results reveal how a continuous, flexible body can transform sparse environmental features into a robust, fault-tolerant network. This mechanism provides a biomechanical framework for understanding the repeated evolution of limbless climbing and offers physical principles for designing agile robots for unstructured terrain.

## Introduction

Arboreal habitats impose extreme physical challenges, where animals must negotiate contacts with branches and tree trunks of variable texture, size, shape, and flexibility, where failure can lead to catastrophic falls. In limbed animals, surface contacts are often established with adapted traits including claws, adhesive pads, grasping limbs, and prehensile tails to meet these demands [1–7]. Snakes, however, present a striking contrast. Despite the lack of limbs and specialized attachment structures, climbing has evolved repeatedly in snakes [8]. Many species are capable of traversing inclined or vertical environments, ranging from narrow branches and dense vegetation to tree trunks, rock faces, and walls [9–13], where contact geometry and availability vary dramatically. How a continuous, limbless body generates support, stability, and propulsion across such diverse terrain remains poorly understood.

Previous studies have primarily examined environments that allow snakes to wrap around a substrate, brace between opposing surfaces, or push against regularly distributed obstacles. On thin branches and tree trunks, snakes commonly climb upward using concertina locomotion, involving cyclic anchoring and advancing body segments, and constrict the surface to support their weight [9– 13]. With anchored regions supporting the animal’s weight, other regions advance or extend across gaps while maintaining posture [14–17]. In confined, inclined channels, snakes instead brace against opposing walls to increase normal forces and thereby generate the friction needed for support [18]. Surface protrusions on poles or within channels allow a shift in gait to lateral undulation, involving waves propagating down the body, by providing discrete contact sites to push against. Lateral undulation is favorable in that it is substantially faster [10, 19] and presumed to be more energetically efficient than concertina [20–22].

Many surfaces encountered by snakes are flat or nearly flat such as large tree trunks, rocky surfaces, and brick walls where constriction and bracing between opposing sides becomes impossible. That snakes climb such surfaces at all therefore poses a basic mechanical puzzle: how can a limbless body use discrete, sometimes sparse asperities to support, stabilize, and propel itself during vertical ascent and descent? Friction remains essential, and some species exhibit remarkable control over their ventral scales, flaring them and forming keels that grip or wedge against surface features [18, 19]. Yet not all species can climb [23, 24], indicating that the limbless body plan provides an opportunity rather than a guarantee.

Understanding how snakes exploit discrete surface features therefore requires treating climbing as a dynamic multi-contact problem. Unlike limbed animals, whose contacts are generally associated with a fixed set of appendages, snakes can establish contacts at many locations along a continuous, deformable body. In limbless locomotion, cycles of limb placement are replaced by transient body–substrate connections that form and propagate along the body as it deforms [25], allowing forces to be applied at multiple locations simultaneously. This distributed contact network provides substantial mechanical redundancy, but how snakes redistribute forces as contacts are gained, lost, and repositioned remains largely unknown.

In this study, we recorded climbing behavior in cornsnakes (*Pantherophis guttatus*) ascending and descending a flat, vertical wall instrumented with force sensitive posts, combining kinematic marker tracking data with time-resolved measurements of the forces at each contact. We found climbing to be quasi-static: net force and torque remained close to balance, and accelerations were small throughout each trial. The contact network was also highly redundant, with snakes maintaining far more contacts than planar force and torque balance requires. A minimal computational model showed that a prescribed body wave with entirely passive contact mechanics was sufficient for climbing, and an open-loop-controlled robot demonstrated the physical viability of this strategy. Snake descents closely matched this passive prediction and were dominated by dissipative contacts. During ascent, however, snakes actively redirected contact forces beyond the predictions for passive Coulomb friction. Although overall force patterns changed when posts were shortened or spaced further apart, snakes used similar stereotyped, balance-preserving force rearrangements across conditions. Together, these results reveal how a continuous, limbless body can climb surfaces that cannot be continuously wrapped or gripped, while providing a framework for comparing climbing strategies across substrates and designing limbless robots for vertical, complex terrain.

## Results and Discussion

To study limbless climbing in a regime where the body cannot wrap around the substrate, we built a smooth acrylic vertical climbing wall with a single column of force-sensing posts^1^(see Methods). Figure 1a-c shows our apparatus, and Fig. 1d shows a sequence of postures and associated force configurations during a typical ascent. Many simultaneous contacts were distributed along the body with forces generally pointed diagonally downward. The corresponding body-curvature kymograph (Fig. 1e) shows a head-to-tail propagation of lateral body bends that was typical for this wall configuration. Horizontal and vertical forces were measured (in the plane of the wall) on each post through time (Fig. 1f,g). Using both time-resolved local kinematics and forces, we computed the net torque on the snake body about the center of mass through time (Fig. 1h).

**Figure 1:**
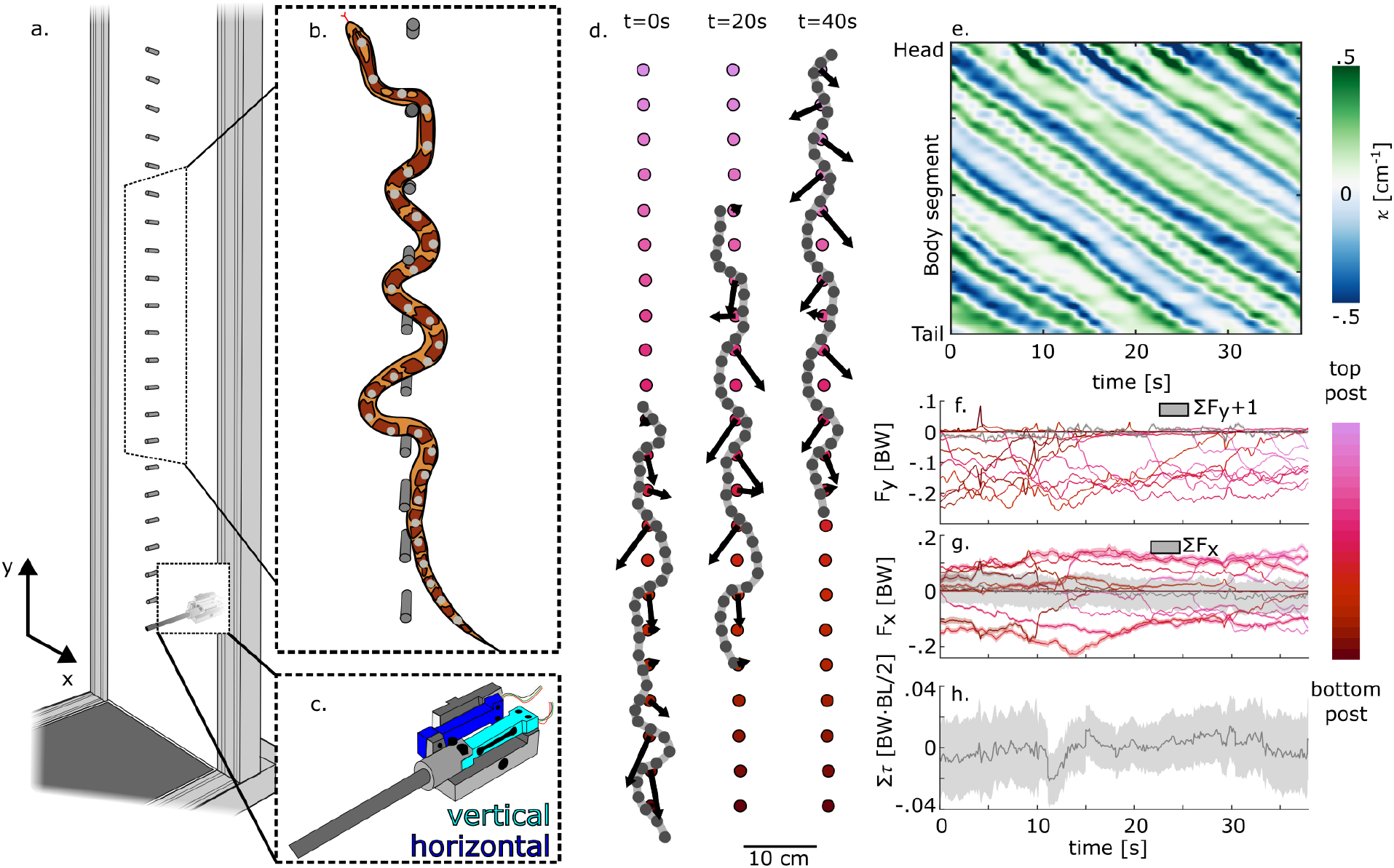
A force-sensing wall resolves body kinematics and per-post forces throughout climbing. **(a)** A single column of 22 force-sensing posts protrudes through a smooth acrylic wall. **(b)** A cornsnake (*Pantherophis guttatus*) climbs by weaving through the posts; reflective markers on its back are tracked by an IR camera array. **(c)** Each post couples to two load cells measuring vertical (y) and horizontal (x) force. **(d)** Three snapshots from a representative ascent: splined body (gray), markers (dark gray), and per-post forces (arrows); posts here are 25 mm long, spaced 50 mm apart. **(e)** Body-curvature kymograph (body position vs. time, colored by curvature κ); lateral bends propagate head to tail. **(f**,**g)** Vertical **(f)** and horizontal **(g)** force on each post over time; summed vertical force tracks body weight and summed horizontal force stays near zero. **(h)** Net torque about the center of mass is centered around zero.

Cornsnakes were chosen as the focal species for this study as they are adept climbers, and their musculature and movements have been studied extensively in other contexts [9,10,19, 26]. Five cornsnakes were used in this study under three post configurations on the wall: 2.5-cm posts spaced 5 cm apart, 0.3-cm posts spaced 5 cm apart, and 2.5-cm posts spaced 10 cm apart. Each snake managed ascent and descent in all conditions (SI section S2) and Fig. 2 summarizes their performance.

**Figure 2:**
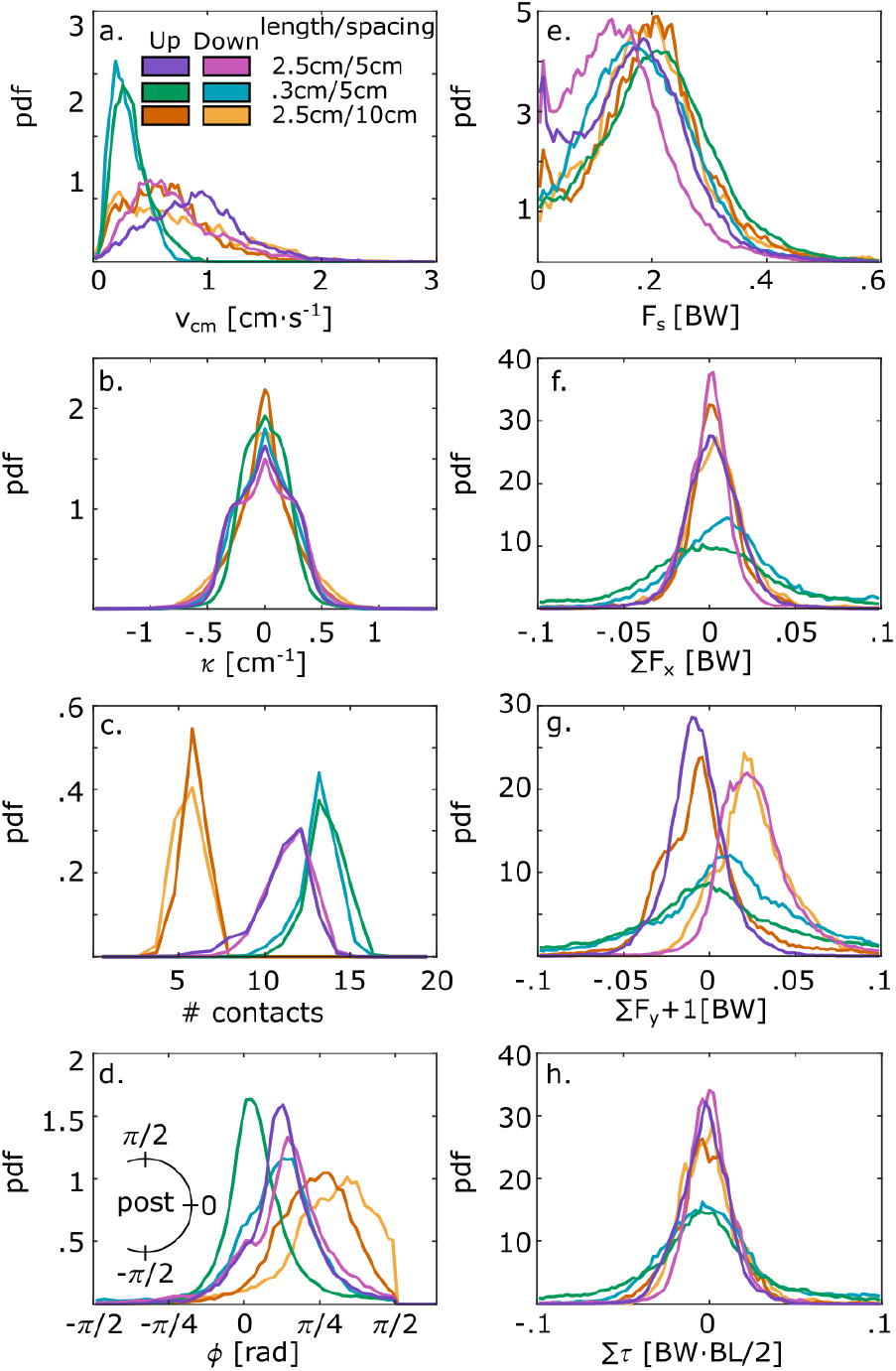
Across substrates, snakes have variable kinematics but maintain force balance. All panels are probability densities pooled across trials, for ascent and descent in three post configurations (legend, a). **(a)** Center-of-mass speed: similar up and down, slower on shorter posts. **(b)** Body curvature κ: similar across conditions. **(c)** Number of simultaneous contacts: more on closely spaced posts, fewer when widely spaced. **(d)** Radial contact location on each post, which shifts with condition. **(e)** Total force per post: similar magnitude across conditions. **(f)** Net horizontal force, net vertical force **(g)**, and torque about the center of mass **(h)** all center near balance; variability grows on shorter posts, where the force pattern is more erratic.

For a given post configuration, speeds of ascent and descent were similar (Fig. 2a). On long posts, snakes used lateral undulation and climbed fastest (5–cm spacing Upward: SI Movie S1, 5–cm spacing Downward: SI Movie S2, 10–cm spacing Upward: SI Movie S3, 10–cm spacing Downward: SI Movie S4); for short posts, snakes used a mix of lateral undulation and concertina gaits (Upward: SI Movie S5, Downward: SI Movie S6), which resulted in slower speeds. This gait change is typical of snake behavior when surface features are reduced or absent [10,18,19]. Though several different complicated shapes were used in each trial and the body remained slightly straighter with short posts or wide spacing, distributions of body curvatures were similar across conditions (Fig. 2b). Snakes formed many simultaneous contacts distributed along the body; however, the number of contacts varied with wall configuration: for closely spaced posts, they typically engaged 8–16 contacts and contacted slightly more with shorter posts; with a wider spacing they had fewer contact points to work with and successfully climbed with 5–7 contacts (Fig. 2c).

Radial contact location on the post varied across conditions. Snakes typically contacted the posts 30–45^°^ above horizontal, but during ascent, shortening the posts shifted contact locations toward the sides of the posts, whereas increasing post spacing shifted contacts toward the tops of the posts (Fig. 2d). Because force direction depended strongly on contact location, these shifts produced more lateral forces on shorter posts and more vertically aligned forces at wider spacing. Force magnitude per contact remained similar across conditions (Fig. 2e). However, due to an increased number of contacts, a greater total force was exerted on shorter posts (Fig. 2c) and was directed largely laterally, consistent with additional bracing on the more challenging substrate (SI section S3, Figure S2). By contrast, wider spacing reduced the number of contacts, requiring the remaining forces to align more vertically to maintain support.

### Climbing is quasi-static

Despite marked differences in the kinematics and overall force pattern across conditions, the total force and torque remained nearly balanced at each moment in time for all conditions.

Force directions, which depended on snake posture and contact location, alternated left and right along the body but the net horizontal force summed to zero overall (Fig. 2f). The measured vertical forces in each case were approximately equal to the snake’s body weight (within a few percent, depending on the post configuration), indicating accelerations up and down the wall were small or absent (Fig. 2g). By calculating acceleration directly, we confirmed this was the case (SI Figure S3). We calculated the total torque about the snake’s center of mass using a lever arm to each post and found this also remained small and centered on zero (Fig. 2h). Climbing therefore proceeded quasi-statically: at each instant the body approximately balanced net forces and torques. More formally, the snake satisfied three constraints:

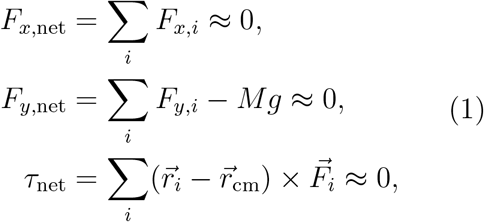

where τ_net_ is the torque about the center of mass and Mg is the animal’s weight. ^2^.

The three constraints in Eq. 1 could in principle be met by as few as three contacts, yet the snakes maintained far more in nearly every condition (Fig. 2c). In this scenario, the snake can access a wide range of force distributions that all satisfy the quasi-static constraints. Even in the short-post condition, where transient departures from balance were more common, the net force and torque distributions remained centered within a few percent of balance, indicating that most of the time, the body nearly maintained quasistatic equilibrium. However, global force and torque balance do not uniquely determine how individual contacts contribute to locomotion. In particular, they do not reveal whether contacts act to oppose motion, as in passive friction, or whether they contribute to forward progression. Therefore, to determine the minimal requirements that are mechanically necessary for limbless climbing, we built two complementary models of climbing on a post array.

### Computational and robotic models establish a minimal climbing baseline

To understand how snakes utilize surface feature contacts, we first constructed a quasistatic computational model of a climbing snake interacting with posts in its environment. The the snake body is represented as a chain of connected segments undergoing a prescribed serpenoid wave (see Methods). As the body moves through the array of posts, each segment experiences up to three environmental forces. At posts, an effective spring force normal to the body enforces local contact geometry and kinetic Coulomb friction opposes local motion across the contact, while gravity acts on all segments (Fig. 3a,b). At each instant, the whole-body translation was determined by enforcing net force and torque balance (see Methods). Surprisingly, this simplified model produces successful climbing behavior both up and down establishing an important minimal baseline: passive contact forces are sufficient to produce quasi-static climbing.

**Figure 3:**
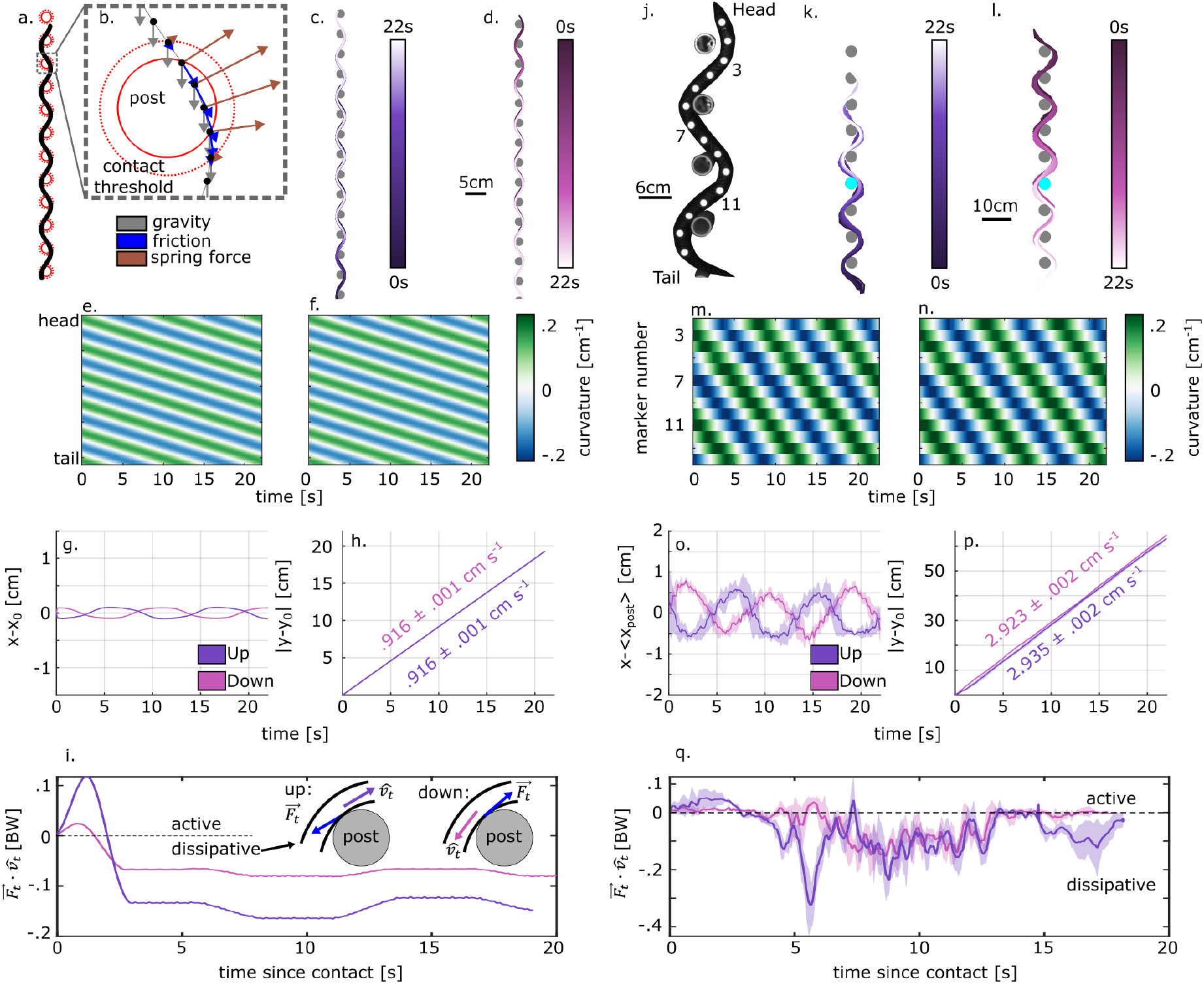
Computational and robotic models reveal the minimal requirements for climbing. *(a–i) Computational model*. **(a)** A chain of segments propagates a prescribed serpenoid wave through the post array; **(b)** each contacted segment feels gravity (gray), Coulomb friction (blue), and a repulsive spring force (brown). The model climbs both up **(c)** and down **(d)**, cycling through serpenoid shapes as bends propagate head to tail identically in both cases. (curvature kymographs: **e**, up; **f**, down). **(g)** Lateral and **(h)** vertical center-of-mass displacement show negligible net horizontal motion over a cycle and constant-velocity vertical advance—consistent with quasi-static climbing. **(i)** The tangential force at a single contact opposite the tangential velocity showing dissipation after a brief adjustment period. Note: here all contacts behave the same. Inset: for dissipative contacts, tangential force opposes local velocity. *(j–q) Open-loop-controlled robot*. **(j)** A chain of servo motors propagates a serpenoid wave, with reflective markers tracked at each joint. The robot climbs up **(k)** and down **(l)** using only 3–4 contacts, far fewer than the snake. Cyan dots in **(k**,**l)** represent the location of the force-sensitive post. Curvature kymographs (**m**, up; **n**, down) and center-of-mass displacement (**o**, lateral; **p**, vertical) again indicate constant-velocity, quasi-static climbing. **(q)** The average tangential force against tangential velocity at the force-sensitive contact again showing dissipation after a brief adjustment.

The success of passive contact dynamics in our computational model inspired us to design a robotic model to confirm the viability of this scheme (Fig. 3j). Because climbing is quasi-static, balance requires satisfying only the three constraints listed in Eq. 1, which provides a theoretical minimum number of contacts needed By propagating a serpenoid wave down the body, the robot was able to climb up and down a vertical post array with no environmental sensing or mechanical force feedback (Fig. 3k,l) (Upward: SI Movie S7, Downward: SI Movie S8). The robot maintained only 3–4 contacts at a time, many fewer than the snake, confirming a near-minimal amount of contacts required for successful behavior. The curvature waves were identical in the two directions (Fig. 3m,n), so climbing direction required no change in gait (see Methods). The robot’s kinematics were also consistent with a quasi-static system as the horizontal center of mass was centered around zero (Fig. 3o) and the vertical center of mass advanced at nearly constant velocity (Fig. 3p), indicating little to no acceleration. Though our force sensors were not able to support the weight of the robot, we designed a wall where one post was equipped with a 6-axis force/torque sensor to measure forces exerted by the robot throughout each climb. To ensure that we captured force dynamics throughout a climb, we mounted this sensor to a middle post (fourth from the bottom) and measured the forces before, during, and after the robot interacted with that post. After a brief adjustment period following initial contact, interactions with this post were generally dissipative, confirming that climbing is physically allowed with friction dominating at contacts (Fig. 3q).

The simulation and robot are complementary: the computational model, with friction-only contacts, shows that a propagating body wave and passive friction are sufficient in principle, while the robot confirms this physically and with only a few contacts. Together they establish a minimal baseline—open-loop body-wave propagation with passive, frictional contacts is enough to climb quasi-statically, requiring neither the snake’s redundant contacts nor feedback.

### Snakes actively redirect forces during ascents

To test whether snakes use this minimal strategy, we calculated the energy transfer direction at each contact. Sliding contacts were considered *dynamic* where the local velocity of the body exceeded 3 mm/s. If the local velocity did not meet this threshold, the contact was considered *static*. For each dynamic contact, we decomposed the measured force and local velocity into components tangential and normal to the average radial contact location *ϕ*. Given the extended nature of the snake body, we used a distance-weighted average of angular contact locations to define a contact centroid; a similar approach was used to estimate contact sliding velocity (SI section S5). We define the tangential contact power to be *P*_*t*_ *= F*_*t*_ · *v*_*t*_, where *F*_*t*_ and *v*_*t*_ are the force and velocity components tangential to contact location defined by *ϕ. P*_*t*_ is negative when the tangential force opposes local sliding direction, indicating a *dissipative* frictional interaction; *P*_*t*_ is positive when tangential force aligns with the sliding direction— and does positive work on the body—which passive Coulomb friction cannot.

At most contacts during experimental descents, the snake’s tangential velocity sliding down the wall was opposed by a tangential force on the snake directed up the wall (*P*_*t*_ < 0; Fig. 4a), consistent with the dissipative interactions predicted by the simulation and robot. Assuming these dissipative contacts are purely frictional gives an effective kinetic friction coefficient *µ* ≈0.22. Upward climbs, however, exhibited reversed dynamics: at most contacts, the snake generated a tangential force differing from the passive-friction prediction by roughly 2*µF*_*n*_, overcoming the resisting friction and driving an equal force in the opposite direction to do positive work on the body.

**Figure 4:**
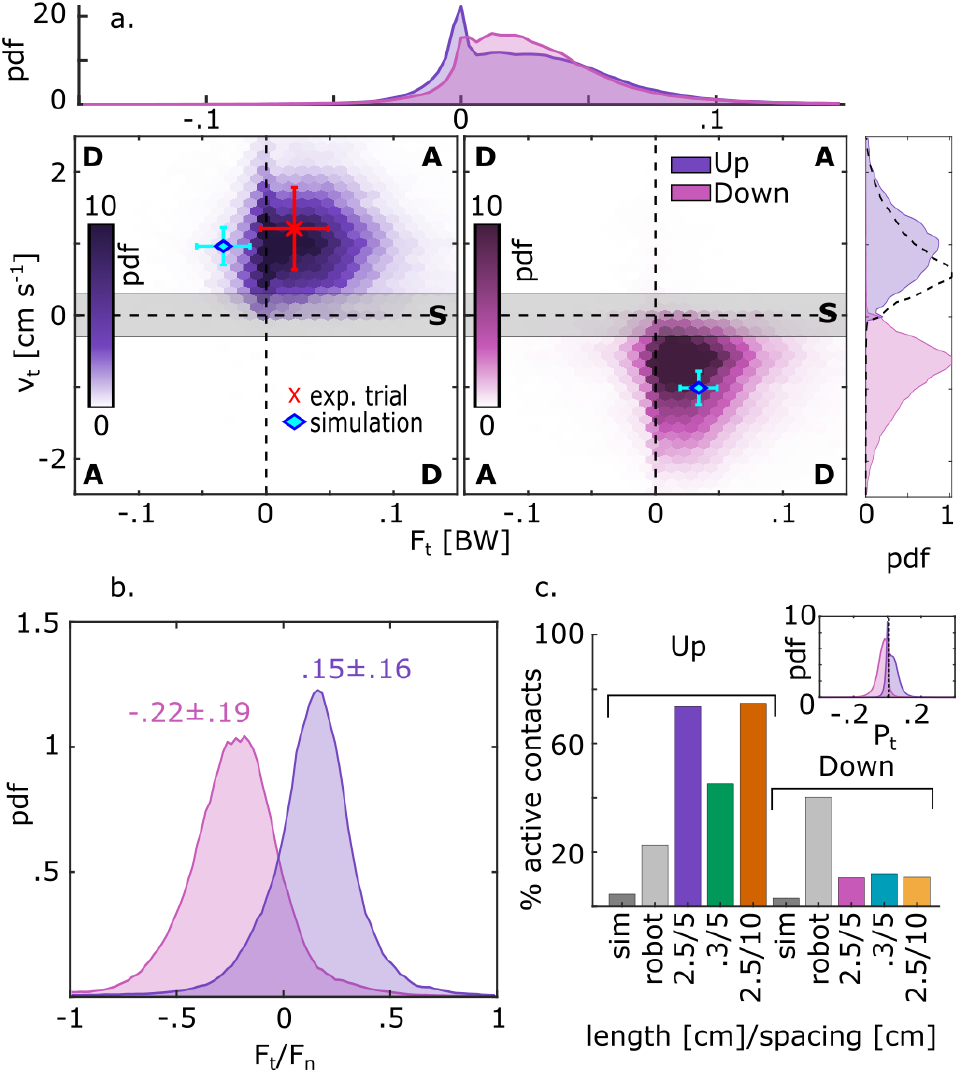
Downward climbs have mostly dissipative interactions at contacts while upward climbs are active. **(a)** Heatmaps of F_*t*_ and v_*t*_ with quadrants marked “A” and “D” respectively indicate active and dissipative contacts respectfully. Dissipation in downward climbs matches simulated predictions while upward climbs, including the trial that the model was based on (red “x”) involve active contacts. Distributions of tangential forces and velocities are shown atop and to the right of the main plots, respectively.**(b)** The ratio of tangential and normal forces in downward climbs matches closely with the µ predicted from dissipative contacts. In contrast, this ratio is nearly opposite in upward climbs. **(c)** The percentage of active contacts in each condition is much higher for upward climbs while downward climbs lie closer to model predictions.

We refer to these positive-power contacts as *active*; meaning that the measured tangential force does positive mechanical work on the body relative to local sliding (*P*_*t*_ > 0). Critically, *P*_*t*_ > 0 is not a generic consequence of propagating a body wave through the array. Our open-loop-controlled robot, which does exactly that with no sensing or control, produces dissipative contacts on average (SI Figure S7). Active contacts therefore reflect an animal-specific contribution beyond this minimal feed-forward mechanism. We stress that “active” remains an energetic statement, not a claim about neural control: our measurements cannot distinguish whether the animal redirects force through sensory feedback, a feed-forward muscular program, or the passive mechanics of a deformable body cross-section. More than 40% of dynamic contacts were active during ascent in every environmental condition, whereas active contacts were much less common during descent, which more closely matched simulation predictions (Fig. 4c). Shortening the posts modestly reduced the prevalence of active contacts, consistent with the shift toward a more concertina-like gait with a greater fraction of static contacts. The robot-post interaction was also occasionally active, but only briefly, when stick–slip dynamics deflected the contact force in the direction of local motion.

The energetic contrast between ascent and descent arose primarily from the reversal of sliding direction rather than from a change in tangential force. Tangential-force distributions were similar for both climbing directions, whereas the tangential-velocity distributions nearly coincided after a sign reversal (Fig. 4a, dashed black line). Consequently, similarly directed forces tended to oppose sliding during descent but align with sliding during ascent, shifting contact power from predominantly negative to frequently positive. Because both the simulation and robot could climb upward using mainly dissipative contacts, this positive work was not required for ascent itself. Instead, it points to an additional force-redistribution strategy used by the animals, raising two questions: how do snakes generate these active interactions, and what functional advantage might they provide?

### New contacts trigger stereotyped, balance-preserving force redistribution

The contact-power analysis describes the mechanical role of each contact in isolation, but climbing stability depends on how the snake coordinates several contact interactions at once. With many redundant contacts, several different force configurations 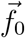 satisfy force and torque balance and make up a “balance space” **B** (Fig. 5a). This wide range of solutions allows the animal to change the load on any individual post without necessarily departing from force balance. We therefore asked how snakes redistribute forces across the contact network each time a new post is engaged.

**Figure 5:**
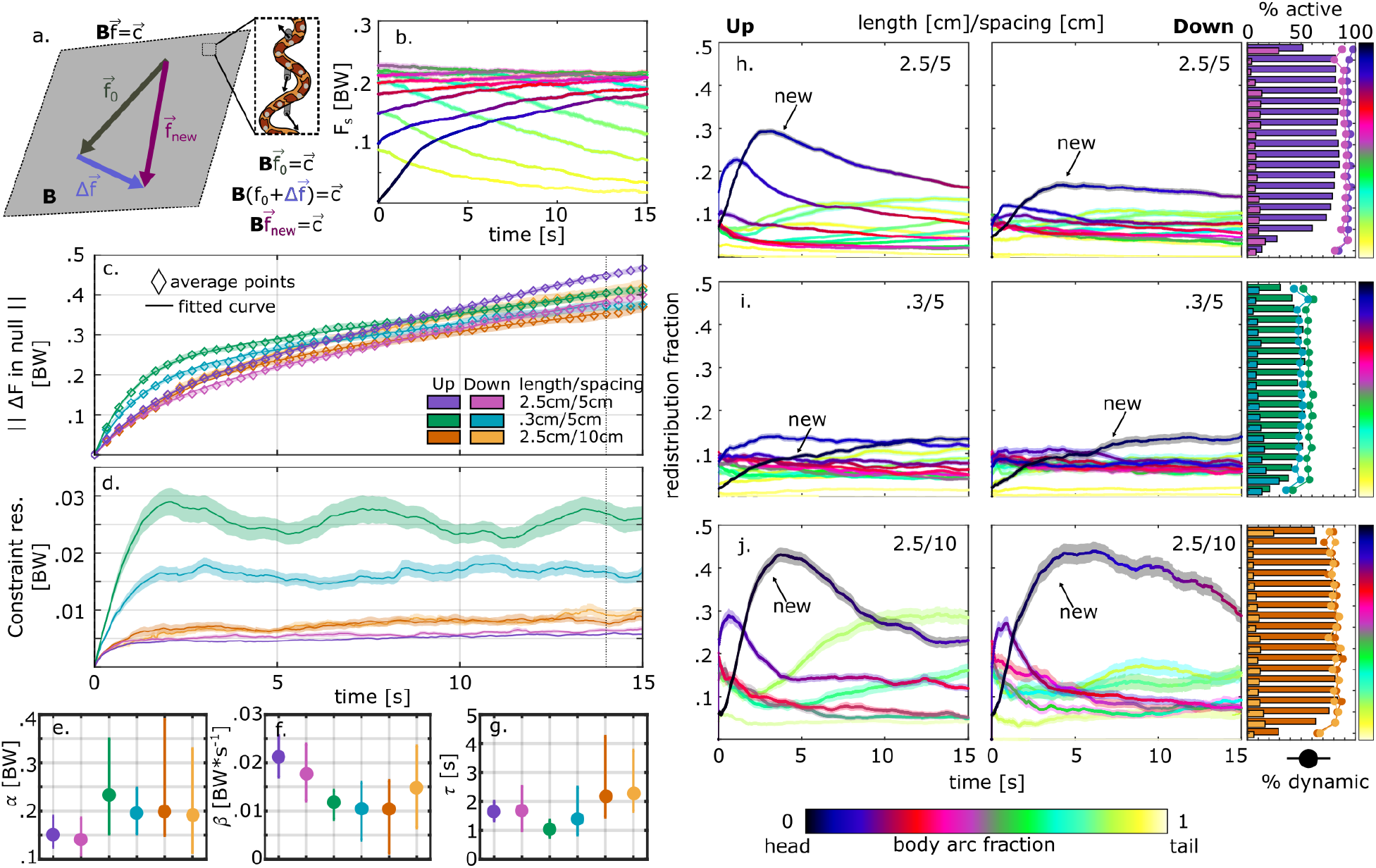
Force redistribution is stereotyped and balance-preserving. **(a)** The balance constraints 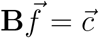 define a space of balanced force configurations; a rearrangement 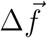 in the nullspace of **B** moves between them 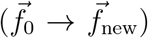 without disturbing balance. Contact-triggered average of the force on each post: force ramps to a stable value on the new contact and its near neighbors, a stereotyped incorporation. **(c)** Magnitude of the balance-preserving redistribution 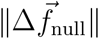 over time (average points and fitted curves), all six conditions. **(d)** Residual force outside the nullspace (the balance violation): it rises and then holds at a small, steady level—largest on short posts, but within a few percent of body weight. **(e–g)** Fitted parameters across conditions: amplitude *α* **(e)**, drift rate *β* **(f)**, and timescale *τ* **(g)**. (h–j) Fractional contribution of each contact to the redistribution in (c), colored by body position (head to tail), climbing up and down; the new contact is labeled. The redistribution concentrates near the new contact on long, closely spaced posts, spreads along the body on short posts, and is dominated by the new contact at wide spacing. The plots on the right track the percentage of contacts along the body that are dynamic and active showing how local force rearrangements affect energetics.

Incorporating a new contact is not an independent action; it triggers a repeatable redistribution of force through the contact network as force ramps up on the new post and eventually reaches a stable, stereotyped value (Fig. 5b). We represent the set of contact forces as a vector 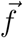 and the quasi-static balance conditions as 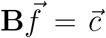, where the three constraints fix the net horizontal force, the net vertical force, and the net torque about the center of mass. Any rearrangement, 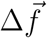, to this force configuration separates into two mechanically distinct parts: a balance-preserving part lying in the nullspace of **B**, leaving the constraints 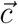 satisfied, and a residual part that violates whole-body balance. The nullspace is precisely the freedom that redundancy provides: within it, the animal can reshape how force is shared among posts without perturbing its equilibrium. We quantify rearrangements across trials by defining t = 0 to coincide with the onset of each new contact and tracked the change in the contact-force vector relative to its value at that instant, focusing first on the balance-preserving (nullspace) component, 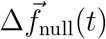 (see Methods, SI section S6).

Across all conditions, the magnitude 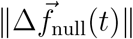 rose rapidly after a contact formed and then continued to drift as the animal advanced through the array (Fig. 5c). We summarized this response with a saturating exponential plus a linear ramp,

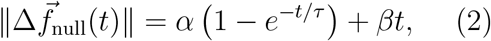

where *α* is the amplitude of the rapid, balance-preserving redistribution recruited by the new contact, *τ* is the characteristic time over which that transient settles, and *β* is the rate of the slower, ongoing reassignment of force as body position, contact geometry, and moment arms change during the climb. We interpret this fit as a description rather than a mechanistic claim: the exponential term captures the transient response to a newly available contact, and the linear term reflects the animal’s continued motion through a changing mechanical environment.

The part of the rearrangement that violated force and torque balance rose over the first second or two and then held at a small, steady level rather than decaying away (Fig. 5d). This also provides a proxy for how closely the system remains quasi-static. With longer posts, less than 1% of the force was outside of balance, however, shorter posts led to more variable force patterns and greater departures from being quasi-static.

Across the six conditions, the transient amplitude α was smallest on long, closely spaced posts and larger when posts were shortened or spaced farther apart (Fig. 5e), though these differences were comparable to the fit uncertainties. The ongoing drift rate β showed an opposite, and clearer, pattern: largest on long, closely spaced posts and roughly half as large on shorter or sparser ones (Fig. 5f), though uncertainties overlapped here as well. The rearrangement timescale τ remained similar in each condition (~1-3 s) (Fig. 5g). Together, these results point to a tradeoff. With dense, reliable contacts, each new contact triggers a small force rearrangement (smaller *α*) but forces must propagate down the body at a higher rate (larger *β*). When the surface features are shorter or sparser, the opposite is required: each new contact requires a large force rearrangement (larger *α*) but loads can be transferred between body segments slower (smaller *β*).

### Body segments are recruited for rearrangement based on the environment

Having established that new contacts trigger mostly balance-preserving redistributions, we next asked which parts of the body contribute most to these force changes. Although redistribution occurs across the contact network, each force is generated at a localized body–post interaction. The spatial pattern of contributions therefore reveals whether contact incorporation is concentrated near the newly formed contact or distributed more broadly along the body.

To investigate, we calculated the fractional redistribution contributed by each contact. On long, closely spaced posts, the established contact just behind the new one initially carried the most rearrangement, but later on the new contact became the dominant contributor to the rearrangement. Middle and hind segment contributions were small and steady until the tail released the trailing post (Fig. 5h). This concentration was strongest during ascent and was responsible for redirecting the tangential force; the region where more dynamic contacts were active occurred just behind the head-most region of the body.

For descent the same trend held, but contributions were more evenly shared along the body while the redistribution maintained dissipative interactions. When posts were shortened, no body segments emerged as relatively strong redistributors (Fig. 5i). Instead, when using concertina-like gaits with multiple sliding and static contacts on the body simultaneously, more body segments were actively involved and contributed equally to the force redistribution. We observed an increase in mid-body adjustments in this environment: on these substrates, the mid-body sometimes flipped over a post, engaged it from the opposite side (Movie S9), and recruited body regions that were previously uninvolved. When posts were spaced farther apart, the new contact made a larger and more persistent contribution, as expected when each contact carries greater weight in a sparser network (Fig. 5j). Together, these results show that each new contact is incorporated through a redistribution of force across the existing network that is largely balance-preserving, but whose size and spatial spread depend on the density and reliability of the available surface features. The limbless body thus uses redundancy not only as excess support, but as a mechanical space in which local forces can be reorganized while maintaining stability.

## Conclusions

For an elongated, limbless body, generating stabilizing and propulsive force on a flat, vertical surface requires much more than generating friction at isolated footholds. Instead, it is important to manage a redundant contact network as a whole to distribute force across contacts as new ones are established. Our computational and robotic models proved that passive environmental contact forces coupled with simple body-wave propagation is sufficient for climbing, but cornsnakes operate in a much richer regime. They engage far more contact points than balance strictly requires, actively redirect force beyond passive friction during ascent, and smoothly incorporate each new contact through a highly coordinated, balance-preserving redistribution.

This gap between what is mechanically sufficient and what the animals do reveals both an opportunity and a demand of the limbless body plan. A continuous body can engage many features at once, opening a redundant space of force distributions where coordinated signals can utilize many different solutions to maintain balance [27]. But this redundancy is not automatically useful: an elongate body does not by itself guarantee climbing. Kingsnakes failed on the same wall (Movie S10). Cornsnakes succeeded by organizing their scattered contacts into a balanced, force-sharing network. With active control localized on segments of the body at each point in time, the exact mechanism by which force is redirected remains unknown. Finely controlled muscles connecting the ribs and backbone to the skin are also highly localized and we posit that these may be responsible for redirecting force [28]. Whether force redirection is neurally controlled or a passive consequence of body deformations falls outside the scope of our data. Whatever the mechanism, doing positive work at a contact rather than merely resisting it re-quires actuation coordinated with the local sliding velocity, implying finely distributed force sensing along the body [29].

Efficient motor control involves conserving energy by ignoring redundant dimensions [27]. In extreme environments, however, energy efficiency may be secondary to robust locomotor performance and stability. Thus, climbing snakes may utilize a system where redundant contacts provide a wide space of solutions to prevent perturbations that may result in falls. The redundancy itself is a principle that reaches well beyond snakes. Whenever a task can be met in more ways than it constrains, e.g., reaching for an object with a multi-jointed arm [30], grasping with many fingers [31], or stabilizing the body over uneven terrain [32], the surplus degrees of freedom span a subspace where activity can vary without affecting the task, which motor systems exploit to absorb perturbations while protecting task-relevant variables [33]. Here, the snake reveals an example of redundancy: a space of internal forces that a multi-point grasp can apply without changing the net wrench on the object [34]. This allows versatility in internal force patterns which they use to hold a climbing body balanced as contacts come and go. That snakes maintain such a network at the cost of many extra contacts reflects a trade-off familiar across biology and engineering—between the economy of a minimal solution and the fault tolerance of a redundant, over-constrained one [35, 36]. Where a single slipped contact can mean a fall, holding whole-body balance to a small, bounded error may be worth the cost, consistent with the tendency of biological systems in high-stakes environments to favor robust redundancy over minimal viability [37].

These results lend themselves to further development of robotic templates which coordinate signals across a network to produce movement [38–41]. Here we have demonstrated the minimum required controls given that the geometry of the propagated wave matches reasonably with the post spacing. Snake-like climbing gaits have been replicated previously allowing these robots to traverse inclined poles [42, 43]. However, our results work toward expanding the repertoire of passible terrain for these devices.

These principles place the repeated evolution of arboreal climbing in snakes within a broader picture of contact-rich locomotion [20, 44]. Low-curvature surfaces cannot be encircled like branches, yet they offer scattered features that a body can recruit. Limbless animals find success in these situations by aggregating features into a malleable, balance-preserving contact network which may be key to managing habitats containing them. By separating the mechanics that make climbing possible from the strategies animals actually use, this work points to the redistribution of force across a redundant contact network—rather than friction at isolated footholds—as a central feature of limbless climbing.

## Materials and Methods

### Study subjects

Five juvenile cornsnakes were used in this study. All individuals were between 16 and 24 months old, weighed between 77 and 114 g, and were between 69 and 82 cm long (SI section S8). The snakes were fed every 7 days. Individuals were eligible to be used for trials on a given day if they: had not eaten in the last 24 hours, were not in shed, and had not been used the previous day. Each individual was not allowed to complete more than 10 trials per day and, in total, each completed between 38 and 54 trials over the whole study amounting to 227 trials in total (SI section S2). All experiments took place in the same room that the snakes were housed. The room temperature was consistently 75-80^°^F and the ambient relative humidity fluctuated from 15-30%. All experiments were performed in accordance with Emory University IACUC protocol 202100179.

### Experimental apparatus

The climbing apparatus used for the experiments consisted of a flat, smooth piece of acrylic, 152 cm (5 ft.) tall by 30 cm (1 ft.) wide. The wall was mounted vertically facing our array of marker tracking cameras. Protruding from the wall was a single column of metal posts, 6 mm in diameter. The posts were covered with a smooth, 3D-printed sleeve 8 mm in diameter. Foam padding was placed at the base of the wall to ensure a soft and safe landing in the event of a fall. Posts were mounted behind the wall and protruded through holes in the acrylic 10 mm in diameter so that the posts did not touch the inside of the holes. Each post was mounted to custom metal brackets which accommodated two TAL221 load cells from Sparkfun with a 500 g load capacity. One load cell was oriented to measure vertical forces on the post while the other was oriented to measure horizontal forces. Each load cell was connected to its own HX711 analog-to-digital signal amplifier which was connected to an Arduino Mega for data collection. There were 22 posts on the wall which required 44 load cells, 44 amplifiers, and 2 Arduino Megas to collect all of the force signals. We were able to collect vertical and horizontal force measurements from all posts simultaneously at ~10 Hz.

We were able to manually move the acrylic wall back and forth to change the length of the posts protruded. In these experiments, posts were mounted 50 mm apart and were set to lengths of 25 mm and 3 mm. The length of the 3D-printed sleeves on each of the posts matched the length of post protruding from the wall.

To record 3D kinematic data, we used an array of 6 Optitrack Primex22 cameras recording at 60 Hz, 5 tracking reflective markers on the back of the snake and 1 recording video. Kinematic data and force data were later synchronized (see Data Analyses below) in MATLAB for analysis.

### Force-sensor calibration

Each load cell is designed with strain gauges to output a signal in mV linearly proportional to the loaded weight on the post (measured initially in grams). Each load cell was calibrated independently by hanging a series of known masses from each post and calculating the slope of the output signal. Horizontal load cells were loaded horizontally with the help of a pulley system. Because each pair of load cells were coupled, we found cross-signals between the two, i.e., a purely vertical load would affect the horizontal load cell’s reading. To account for this, vertical and horizontal forces were each calibrated using both load cells and both the linear and quadratic responses to the series of static loads on each (SI section S9).

### Experimental procedure

Each day, 2– 3 snakes were made available for experiments to perform upward and downward climbs given that they met the criteria mentioned above. Before performing the experiments, the snakes were marked with 30–40 markers 6 mm (1/4 in) in diameter cut from silver reflective tape and adhered at roughly even intervals down each snake’s back. A trial order was randomly generated dictating the order to use each individual and climb direction.

Before performing trials on a given day, the ends of the posts and four corners of the wall were marked with the same markers as the snakes and recorded with the cameras for 3 s. From this we were able to show the snake position relative to the post positions and the wall position. Markers on the wall and posts were removed before experiments took place. The wall and posts were kept in the same position throughout the day.

A trial started when the individual snake is introduced to the wall (at the bottom of the post column for upward climbs, at the top for downward). Once the snake climbed fully up or down the column of posts, the trial was completed. If the desired behavior was not observed, the snake was repositioned for another attempt. If the snake remained stationary or paused during climbing, we tapped their tail to elicit an escape response. If after 10 minutes no climbing behavior was observed, the trial was skipped and the next random trial in the order was performed. Snakes rested while other individuals performed trials. If a snake was scheduled for trials back-to-back, no rest was allowed if the trial lasted less than 2 minutes. For longer trials, the individual was allowed to rest for at least 10 minutes in between trials. Individuals were subjected to no more than 10 trials and/or 10-minute periods with no desired behavior per day.

### Quasi-static climbing model

To ask what is minimally required for a limbless body to climb through a sparse set of contacts, we built a simple model of a snake weaving between posts [LINK TO CODE]. We represented the body as a chain of *N* = 220 rigid segments of equal mass *m* = *M/N*, where *M* is the total body mass. Each segment had a length *ℓ*_*s*_ = L/(N 1), and the total body length was *L*. Rather than solving for the body shape, we prescribed it as a traveling serpenoid wave—a posture in which the curvature, *κ*(*s, t*), varies sinusoidally along the body and travels from head to tail at constant speed, *v*_*c*_. The curvature is given by

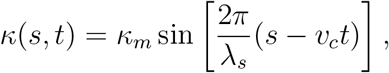

where s is arc-length position along the body, *t* is time, *κ*_*m*_ is the peak curvature, *λ*_*s*_ = *L/n*_*w*_ is the wavelength, given by the body length divided by the number of waves *n*_*w*_. Integrating the curvature along the body gives the segment tangent angles in the laboratory frame,

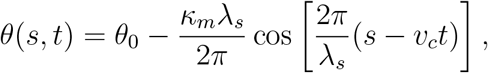

where *θ*_*0*_ orients the body along the direction of climbing. We chose the waveform parameters to approximate the body shapes we observed during climbing, and we set *v*_*c*_ so that the resulting center-of-mass speed matched our measurements.

Every segment experienced gravity, 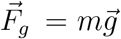. A segment experienced additional forces only when it overlapped with a post. Specifically, when the penetration depth *δ* (the distance by which the segment intrudes past the post’s effective radius) exceeded zero, the segment experienced a contact reaction force and friction; a segment that was not in contact (*δ* = 0) experienced only gravity. We resolved these contact forces in the body frame, using the unit vectors normal 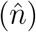 and tangent 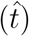 to the segment. The reaction force was a repulsive, one-sided penalty force that resisted interpenetration, given by

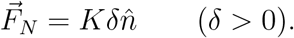

This force grew linearly with the overlap δ (an exponent of one is appropriate for cylindrical contact), with stiffness K, and pointed along the segment normal so as to push the body out of the post. Kinetic Coulomb friction acted along the body and opposed sliding,

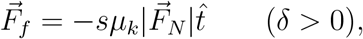

where *µ*_*k*_ is the coefficient of kinetic friction, the normal load is the reaction force magnitude 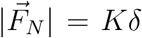, and *s* = sgn(*v*_∥_) is the direction in which the segment slides along its own axis. In other words, the perpendicular direction captured the contact reaction force and the parallel direction captured friction. Gravity was the only force with components along both the parallel and perpendicular directions.

A rigid post can react only along its own surface normal, which coincides with the segment normal 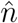 when the body lies tangent to the post, as it does for the contacts we observed. We deliberately apply the reaction along 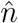 rather than along the exact post normal: directing it along the post normal would couple the force direction to the center-of-mass displacement we solve for at each step, which generally prevents the body from reaching exact force balance for a prescribed shape, whereas the body-normal form leaves the force directions fixed by the posture and admits an exact quasi-static balance.

Consistent with our observation that climbing snakes accelerate very little (Fig. 2), we treated the body as quasi-static, requiring the net force and net torque to vanish at every instant,

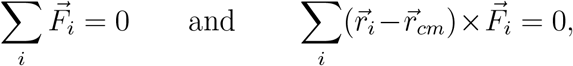

where 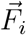 is the total force on segment *i* and 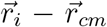 is its position relative to the center of mass. At each time step we held the prescribed shape fixed and solved for the small rigid-body displacement of the center of mass—two translations and one rotation— that brings the net force and torque closest to zero, normalizing the torque by the body length L so that it scales comparably with the forces. The waveform parameters reported in the main text are those that best reproduced the body shapes we observed.

### Robotic model

To test the viability of passive contact mechanics and near-minimal contacts in a physical system, we designed an open-loop-controlled robotic model. The robot consists of 15 servo motors linked together with 3D-printed brackets to create a motor chain with a joint between each motor pair. We sinusoidally varied the joint angles to generate a serpenoid wave. With 2 waves on the body and joint angles between 135^°^ and 180^°^, the robot was able to climb up and down a vertical post array with posts made out of 41-mm-diameter PVC pipes spaced 127 mm apart. One post in the center of the array was connected to an ATI Mini-40 6-axis force/torque load cell to measure forces at a single contact throughout the entire robot-post interaction. Waveform parameters were chosen so that the robot had only 3–4 contacts throughout each trial.

### Data analyses

All data were analyzed using MATLAB. Data and code can be found here: https://osf.io/x5rdu/overview?view_only=3a8c024555c747d0a15d5f2bab60ec9e. Force data were synchronized to points in the marker tracking data using timestamps on each force point and a timestamp for the start of the marker tracking data. Because the force data was recorded at a lower frequency than the cameras, the data points were linearly interpolated to match the 60 Hz kinematic data. Force data were calibrated using calibration matrices for each individual load cell and normalized by the weight of the particular snake.

During climbing assays, the cameras recorded the 3D positions of the 30–40 markers along the back of the snake. Most markers were tracked for the entire duration of each trial, but occasionally some were dropped and reappeared. These markers were re-linked by finding two markers that disappear/reappear closest to the same spatial location. Any trials where markers were dropped and unable to be re-linked were truncated to the longest time span where all markers were present. If the longest time span was greater than 20 s, the trial was excluded from analyses. Once linked through time, markers were ordered by proximity along the body, starting with the one at the greatest height.

We used the position of the wall to transform all of our spatial data into a consistent coordinate system. The markers recorded at corners of the wall were used to fit a plane to the wall. By building an orthonormal basis matrix to describe the plane, we linearly transformed all of the snake, post, and wall marker points relative to this plane. As a result, the plane of the wall became the *xy* plane where *y* is directed vertically and *x* is directed horizontally. The *z*-direction describes how far away from the wall the markers are (SI Figure S5). We also smoothed the data using a 1D Gaussian filter with a standard deviation *σ* = 20 time points and a radius of 4 · *σ* over time to eliminate small perturbations below the noise threshold of the camera array (~0.2 mm).

We calculated the position of the body segments relative to the posts and examine the body segments used to apply the force detected by each post and their shape at each point in time. We used Modified Akima Interpolation to obtain a set of 200 cubic spline points along the snake’s backbone at each point in time. The curvature of the body was calculated from these spline points through time using the Pratt method to fit circles to local regions of 11 points. From here, we used a distance threshold of 17 mm to identify which body segments were close to which posts. We also used a force threshold of 0.21 g, which we found to be 3x the standard deviation of the sensor readings at rest, to show when a contact was established.

We chose to focus on segments of trials in which snakes were actively making upward or downward progress during climbs. Thus, if the snake stopped, i.e., *v*_*cm*_ < 1 mm/s for at least 2 s, we truncated the trial to the longest time span with no pauses. Lengths of 3 s either before or after a pause were also eliminated to avoid sudden speed ups and decelerations. If the longest time span with no pauses was not at least 20 s, the trial was excluded.

Kinematic quantities at each contact such as the radial contact position *ϕ* and tangential velocity *v*_*t*_, were calculated using a weighted average of the quantity measured at every body segment within the distance threshold of the contact. Weights were assigned based on the distance from each point to the contact threshold (SI section S5).

A balance space was constructed based on possible arrangements of applied forces that satisfied the constraints of force and torque balance. We defined a nullspace of this balance consisting of force rearrangements that did not violate the constraints (SI section S6). Contact triggered average plots show the snake’s average behavior relative to when a new contact is formed (SI section S11). These were used to quantify force rearrangements relative to the nullspace and the fraction of rearrangement attributed to each body segment.

## Author contributions

C.A.R, J.R.M, and J.M.R designed research. C.A.R, M.L., Y.N., G.T., and J.M.R. performed research. C.A.R., M.L., and J.M.R. analyzed data. C.A.R., J.R.M, and J.M.R wrote the paper. The authors have no competing interests to declare.

## Acknowledgments

The authors thank Horace Dale and Lowell Ramsey for assistance in experimental design, and Jessica Tingle, Jake Socha, and Gordon Berman for helpful discussions. C.A.R. was partially supported by NSF PHY-2310741.

## Disclosure of Delegation to Generative AI

The authors declare the use of generative AI in the research and writing process. According to the GAIDeT taxonomy (2025), the following tasks were delegated to GAI tools under full human supervision: Code optimization, Proofreading and editing. The GAI tool used was: ChatGPT 5.5, Claude Opus 4.8. Responsibility for the final manuscript lies entirely with the authors.

## Footnotes

1 In the absence of posts, cornsnakes could not make forward progress on the acrylic surface and slid downward even at shallow inclines (SI section S1), confirming that the wall itself provided insufficient support. With posts present, however, all cornsnakes successfully ascended and descended the wall.

2 Forces into/out-of the wall were measured in 17 experiments: they were small and not associated with displacements in that direction. We therefore conclude that the snake is below the static friction limit. See SI section S4.

## Notes

https://osf.io/x5rdu/overview?view_only=3a8c024555c747d0a15d5f2bab60ec9e

### Competing Interest Statement

The authors have declared no competing interest.

### Summary of Updates

Adding revised supplementary material. Updated caption to Figure S8.

