## SupplementalInformation for "Redundant contacts and force redistribution stabilize limbless vertical climbing"

### Supplementary material for "Redundant contacts and force redistribution stabilize limbless vertical climbing"

Movies can be downloaded from: https://osf.io/x5rdu.

**Supplemental movie S1.**

Upward climbing with 2.5–cm posts spaced 50 cm apart. In all of these movies, the black and white video on the left is a video of the experiment where we track reflective markers placed on the back of the snake. The animation on the right shows the tracked body of the snake and the arrows represent the force on each post. Dotted lines show a path across the post perpendicular to the snake's contact location which makes the deflection apparent. Here forces are generally deflected below this line indicating a tangential force opposite what is predicted by friction. Video is 5x speed.

**Supplemental movie S2.**

Downward climbing with 2.5–cm posts spaced 50 cm apart. Video is 5x speed.

**Supplemental movie S3.**

Upward climbing with 2.5–cm posts spaced 100 cm apart. Video is 5x speed.

**Supplemental movie S4.**

Downward climbing with 2.5–cm posts spaced 100 cm apart. Video is 5x speed.

**Supplemental movie S5.**

Upward climbing with 0.3–cm posts spaced 50 cm apart. Video is 5x speed.

**Supplemental movie S6.**

Downward climbing with 0.3–cm posts spaced 50 cm apart. Video is 5x speed.

**Supplemental movie S7.**

Upward climbing from our robotic model. The force sensitive post is colored cyan in the animation. Video is 5x speed.

**Supplemental movie S8.**

Downward climbing from our robotic model. Video is 5x speed.

**Supplemental movie S9.**

Highlighting an instance in movie S5 where the middle of the body changes sides of the post demonstrating active readjustment in the mid-body. Video is 5x speed.

**Supplemental movie S10.**

Brooks' Kingnakes (*Lampropeltis getula brooksi* perform poorly and fail on our climbing wall. Clips are real speed.

### S1 Acrylic surface with no posts

We performed 12 experiments with 2 individuals (ta and ch listed above) to gauge the greatest incline that would be passible for a snake without dedicated surface features on the acrylic wall. We did this by starting with the wall level (parallel to the ground) and placed the snake on one end. We then elevated that end of the wall by hand until the snake slid down. We placed two reflective markers on the snake, one on the head and one on the mid-body, and used our marker tracking setup to track the snake's position over time. Four markers were placed on each corner of the wall and remained visible through the trial. This allowed us to fit a plane to these markers and calculate the incline angle of the wall at each point in time. Figure S1 shows, for a single trial, the snake's velocity relative to the wall end based on the angle of the wall. For these twelve trials, the incline angle where snakes started to slide was $16 \pm 2°$ demonstrating the impossibility of snakes traversing this surface above this incline without post contacts.

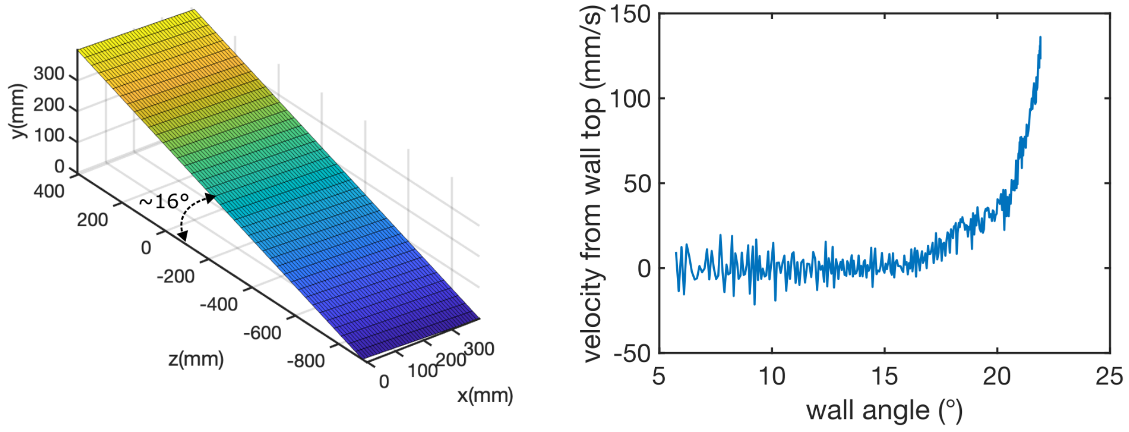


**Figure S1: Wall tilting.** Here we tilted a flat acrylic wall with no posts by hand until it reached an angle (left) where the snake started sliding indicated by its velocity relative to the end of the wall (right).

#### S2 Trial counts by individual

| Animal ID | mass (g) | length (cm) |
|---|---|---|
| ta | 96–103 | 74.4–75.4 |
| ch | 101–112 | 79.0–85.1 |
| el | 99–120 | 71.9–77.5 |
| ca | 80–95 | 73.4–75.2 |
| po | 87–104 | 70.1–73.4 |

**Table S1:** Masses and lengths of the snakes during experimental phase.

| Animal ID | 25/50 | 3/50 | 25/100 | total |
|---|---|---|---|---|
| ta | 10 | 10 | 8 | 28 |
| ch | 11 | 10 | 8 | 29 |
| el | 10 | 8 | 7 | 25 |
| ca | 7 | 9 | 7 | 23 |
| po | 10 | 2 | 9 | 21 |
| total | 48 | 39 | 39 | 126 |

**Table S2:** Number of upward climbs for conditions labeled as post length (mm)/post spacing (mm).

| Animal ID | 25/50 | 3/50 | 25/100 | total |
|---|---|---|---|---|
| ta | 9 | 6 | 3 | 18 |
| ch | 11 | 5 | 9 | 25 |
| el | 8 | 10 | 5 | 23 |
| ca | 10 | 1 | 6 | 17 |
| po | 10 | 2 | 6 | 18 |
| total | 48 | 24 | 29 | 101 |

**Table S3:** Number of downward climbs for conditions labeled as post length (mm)/post spacing (mm).

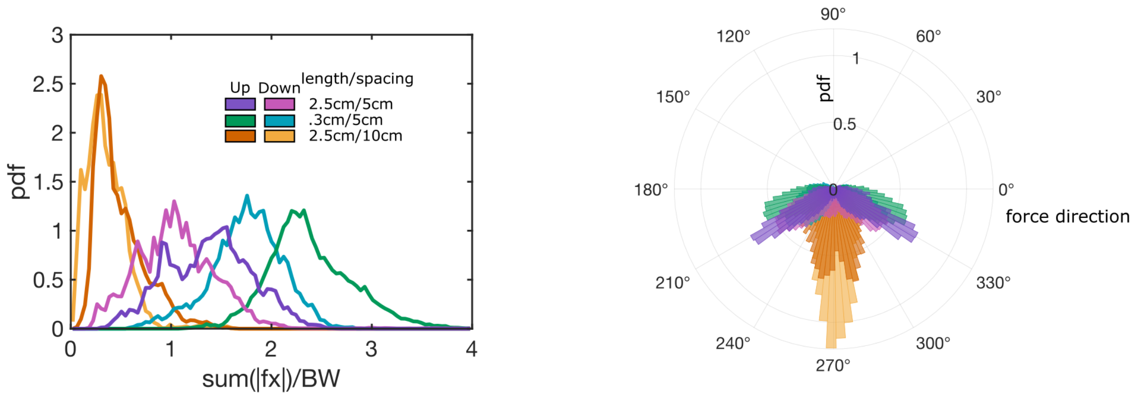


**Figure S2: Bracing force.** (left) The total magnitude of horizontal force at every point in time. (right) a polar histogram showing the direction of force applied to each post.

#### S3 Bracing force

With shorter posts, forces were directed more laterally creating a high bracing effect. While still supporting their weight with an increased number of contacts, lateral forces increased by $\sim 1$ body weight in total magnitude.

#### S4 Forces in the third dimension

A major limitation of our setup is that we can only measure forces in 2D in the plane of the wall. Adding a third dimension of force resolution would be mechanically costly for the setup, but we sought to get an estimation for whether the snakes are pulling on the posts

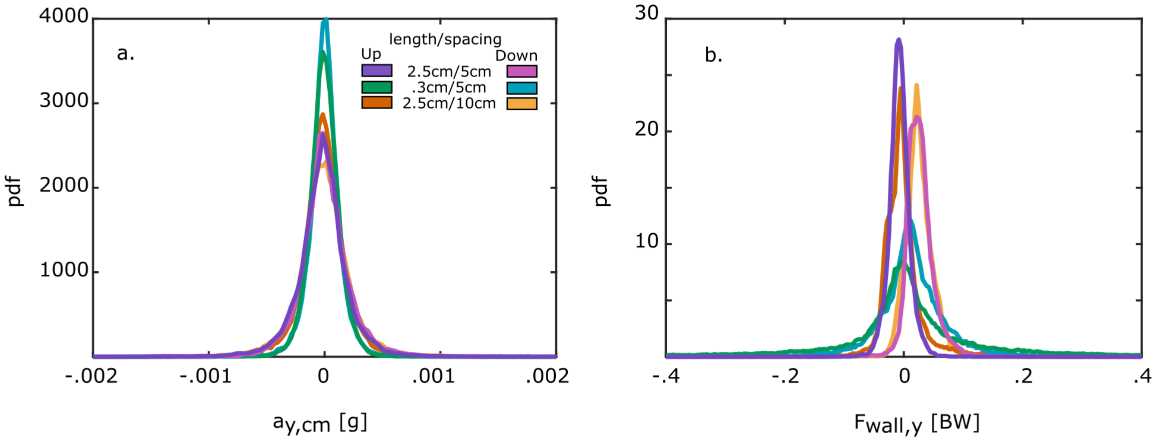


**Figure S3: Accelerations.** (left) The acceleration of the snake's body in each case was either small or zero. (right) Any residual force in the y-direction can be assumed to be from the snake slightly using the wall itself for support.

to generate friction with them and the wall itself. To do this, we replaced the force sensors attached to the 12th post from the bottom with a 6-axis load cell. With this we were able to detect forces directed into and out of the wall at a single point.

We then completed between 3 and 6 trials each with 2.5 cm posts and 0.3 cm posts spaced 5 cm apart; only two individuals (el and ch) were employed for these trials. We found the amount of force that the snakes pull on the posts with not insignificant (Fig. S5). With longer posts, the snake consistently pulled on the post with a force about 1% of its body weight. When the posts were shortened, however, and generating friction became more difficult, the force directed outward on the post increased to over 5% in some instances. This would indicate that there is a friction force directed into the wall on the snake keeping it from sliding outward and cantilevering off the wall.

#### S5 Metrics relative to contact location

To detect individual contacts in our data, the splined snake body must come within a contact threshold of 17 mm and apply a force comfortably outside the noise range of our sensors

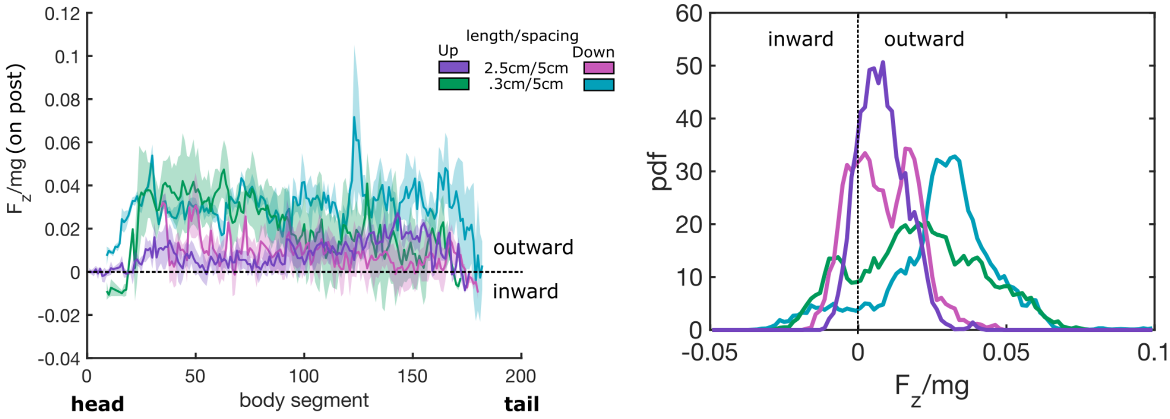


**Figure S4: Third dimension forces directed into and out of the wall.** A six axis load cell allowed us to track if the snakes were pushing or pulling on a single post perpendicular to the wall plane. The left plot shows the average amount of force applied to this post as each body segment slides past. The distribution of pulling force is even along the body except for the very ends. The right plot shows a histogram of all of the third-dimension forces applied to the post which inflate directed outward for shorter posts.

(0.21 g). The radial contact locations $\phi$ and the velocity at each contact using a weighted average of all of the segments within the contact threshold

$$\phi = \sum_{seg} w_{seg} * \phi_{seg}/N_{seg}$$

$$\vec{v} = \sum_{seg} w_{seg} * \vec{v}_{seg}/N_{seg}$$

where the weights $w_{seg}$ are assigned as the distance from the segment to the contact threshold boundary and normalized at each contact and $N_{seg}$ is the number of segments at the contact. $\phi$ was used to break the velocity and force at each contact into their tangential and normal components. Following from this, the tangential velocity was

$$v_t = \sqrt{\|\vec{v}\| - (\vec{v} \cdot \hat{\phi})}$$

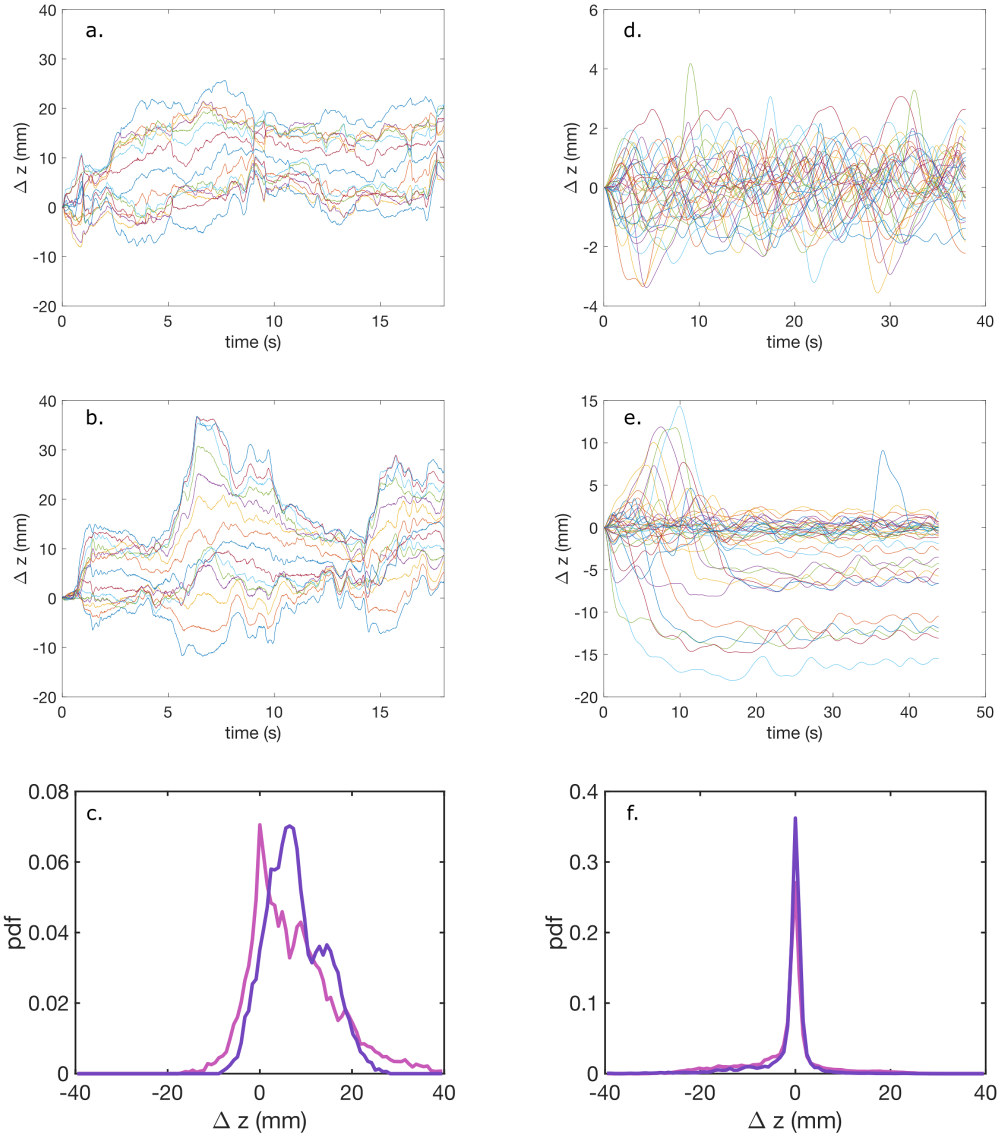


**Figure S5: Third dimension motion.** a-c robot, d-f snake. (a,d) change in z position of each marker over an upward trial. Positive is moving away from the wall, negative is moving toward the wall. (b,e) change in z position of each marker over a downward trial. (c,f) Histograms of the z displacements over all trials in upward (purple) and downward (pink) trials.

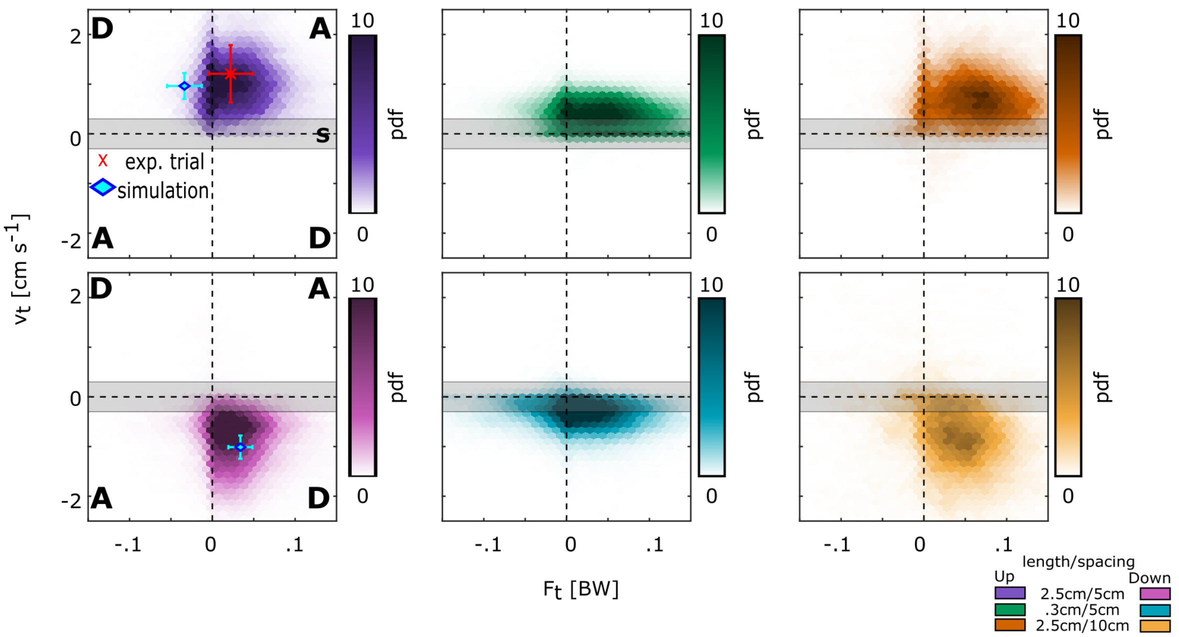


**Figure S6: Heatmaps of $F_t$ and $v_t$ for each post configuration.** Upward climbs in all conditions involve mainly active contacts where downward climbs remain dissipative. There are, however, an increase in static contacts when posts are shortened which is characteristic of the concertina gait.

where $\hat{\phi}$ is a unit vector pointing from the center of the post to the edge at angle $\phi$ and the tangential force was expressed as

$$F_t = \|\vec{F}\| * sin(\alpha)$$

. where $\alpha$, the force deflection angle, is defined as the difference between $\vec{F}$'s direction and an angle 180° from $\phi$. We could then perform the contact analysis displayed in Figure 4 establishing each as active or dissipative. Similar plots for our other conditions can be seen in Figure S6.

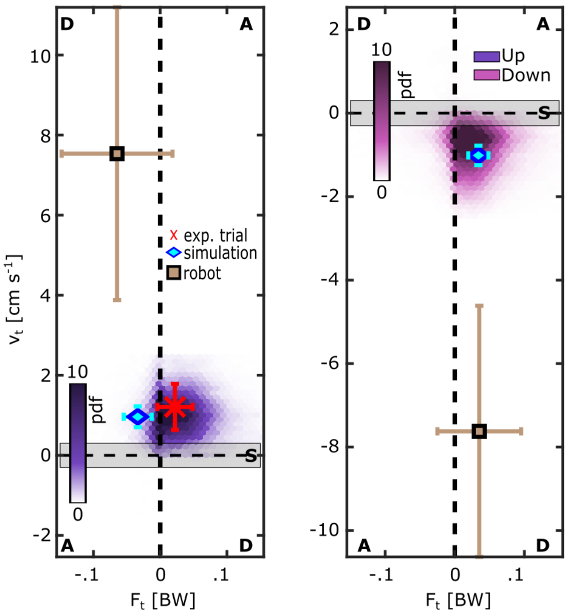


**Figure S7: Heatmaps of $F_t$ and $v_t$ as in Figure 4 including robot measurements.** Contact interactions in robot experiments tend to be dissipative as with the computational model.

#### 78 S6 Nullspace calculation

Each force configuration can be represented by the horizontal and vertical force on each post $\vec{f_0} = [f_{x1}, f_{y1}, f_{x2}, f_{y2}, \ldots, f_{xi}, f_{yi}, \ldots, f_{xN_p}, f_{yN_p}]^T$ where $i$ denotes the post index and $N_p$ is the total number of posts on the wall. The balance space $\mathbf{B}$ can be represented by a matrix

$$\mathbf{B} = \begin{bmatrix} 1 & 0 & 1 & 0 & \ldots & 1 & 0 \\ 0 & 1 & 0 & 1 & \ldots & 0 & 1 \\ -(y_1 - y_{cm}) & x_1 - x_{cm} & -(y_2 - y_{cm}) & x_2 - x_{cm} & \ldots & -(y_{N_p} - y_{cm}) & x_{N_p} - x_{cm} \end{bmatrix}$$

which when multiplied by a valid $\vec{f_0}$ within the space gives $\mathbf{B}\vec{f_0} = \vec{c}$ where $\vec{c} = [0, Mg, 0]^T$ reflecting the constraints

$$\sum F_x = 0$$
$$\sum F_y = Mg$$
$$\sum \tau = 0$$

As the snake establishes new contacts, each change in the force rearrangements must still satisfy the constraints such that $\mathbf{B}(\vec{f_0} + \Delta\vec{f}) = \vec{c}$, thus we can say that the rearrangement $\Delta\vec{f}$ lies within the "nullspace" of $\mathbf{B}$.

### S7 Bootstrapping fit parameters

To find the bounds on the fit parameters for the nullspce function, we used hierarchical bootstrapping. We first resampled the data with replacement for individuals picking 5 each time. We then resampled the trials for each individual, and then resampled events (new post contacts) within each trial. We put bounds on the bootstrapping so that the same number of events was chosen each time. We bootstrapped and fit the nullspace trajectory for 5000 iterations to establish a range for the fit parameters presented.

### S8 Measuring snake length

Without anesthetizing a snake, it is difficult to measure its length directly. Our goal was for these experiments to be as non-invasive as possible so anesthesia was not used. Instead, we measured snake lengths digitally. We first took a picture from directly overhead of the snake resting on a white surface next to a scale bar so that we could assign length values to the image pixels. We then binarized the images in MATLAB, dilated, and eroded the images in order to isolate a silhouette of the snake. We then used a built in skeletonization algorithm to calculate the length of the snake in pixels which was translated to real units with our scale bar.

#### S9 Force sensor calibration

Each force-sensing post on the climbing wall consists of two 500-g capacity load cells. Each load cell contains strain gauges wired in a bridge circuit so that changes in resistance of the strain gauges cause a voltage difference across the bridge corresponding to the applied load. Each load cell is connected to its own HX711 analog-to-digital conversion chip. The digital signals are then read using an Arduino mega.

Each load cell has to be calibrated so that forces can be determined from the raw signals. The raw signals are linearly proportional to the applied load. Independently, the load cells can be calibrated by hanging a series of known masses from the end of the load cell and calculating the slope of the signal produced for each load. We performed a similar series of calibrations for each of our load cells. To calibrate the vertically oriented load cells, we hung weights of 0g, 2g, 5g, 10g, 25g, 50g, 75g, 100g, 200g, 300g, 400g, and 500g from the attached post and recorded the raw signal from each for 10 seconds (Figure S8a). The average of the signal over 10s could be plotted for each load to determine the linear calibration factor that should be used for each (Figure S8b). For horizontally oriented load cells, we used the same sequence of weights, but used a pulley system to load the post directly horizontally (Figure S8a). A series of experiments loading the post at different distances determined that the signal produced was independent of where on the post the load was placed (Figure S8d). All weights in this calibration procedure were loaded 12.7 mm from the end of the post.

The way our load cells were arranged, the signals from each pair ended up being coupled, i.e. a purely vertical load produced a signal read by the horizontally oriented load cell and vice versa (Figure S8b,c).). Therefore, we used a linear system of the cross-signals to reproduce the known load and calculate a calibration matrix to determine measured forces in our experiments from the signals produced by each load cell pair:

$$\overrightarrow{F} = \mathbf{B} * \overrightarrow{V} \tag{S1}$$

$$\begin{pmatrix} F_x \\ F_y \end{pmatrix} = \mathbf{B} * \begin{pmatrix} V_x \\ V_y \\ V_x^2 \\ V_y^2 \end{pmatrix} \tag{S2}$$

where $\mathbf{B}$ is the $2\times4$ calibration matrix, $F_x$ and $F_y$ are the known loads applied horizontally and vertically respectively, and $V_x$ and $V_y$ are the raws signals produced by each load cell ($x$ corresponding to the horizontally oriented one, $y$ corresponding to vertical). Adding quadratic terms to our fitting reduced residuals compared to using linear terms only (Figure S8d).

#### S10 Randomizing trials

Though versatile in the environmental configurations it can replicate, our experimental apparatus requires some level of reconstructing in order to change. We therefore could not randomize environmental conditions as that would require rotating between them. Instead, we completed all of the trials for one condition before moving to the next.

To introduce as much randomness as possible to account for trial ordering, we selected the individuals and climb directions for each trial at random. Each day, 2–3 snakes were made available for trials. For each trial, each snake was assigned a number and a random number generator in MATLAB would select which individual got used. We also had this random number generator choose whether the snake is to climb up or down the wall. We continued with this process until snakes reached their trial maxima for the day. If one snake completed its quota before the others, it was removed from the random pool and trials continued randomly with the remaining snakes.

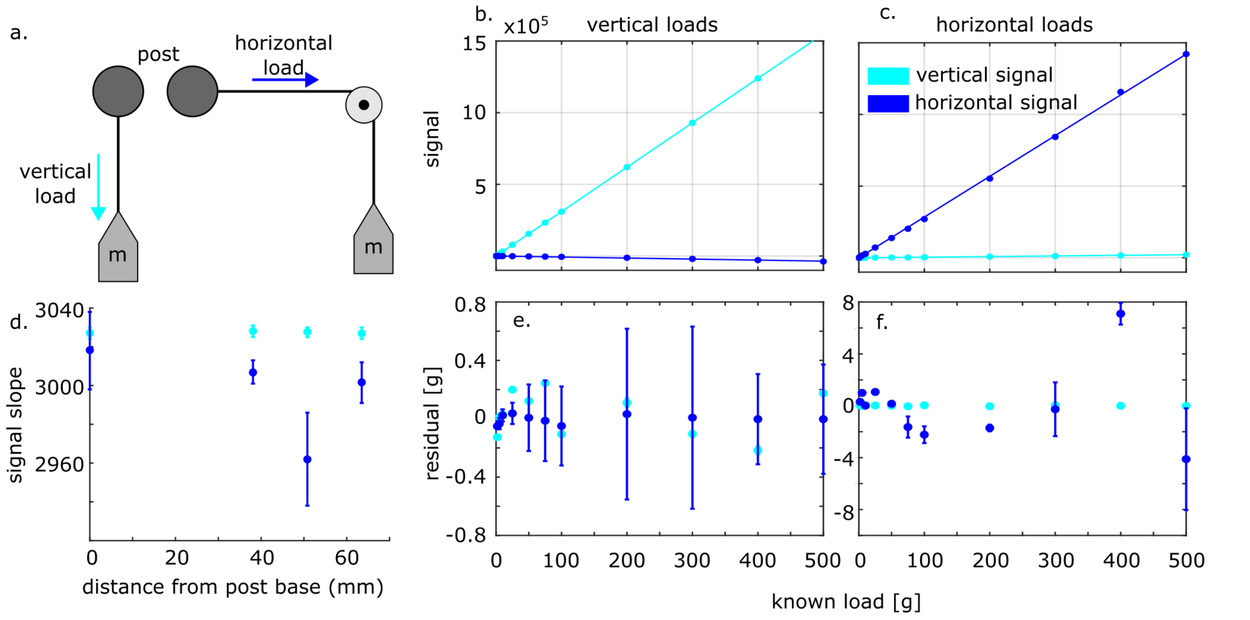


**Figure S8: Force sensor calibration (a)** A series of known weights were applied to each force sensor both vertically and horizontally to calibrate the two load cells. Vertical loads **(b)** and horizontal loads **(c)** caused a large linear signal in the load cell oriented in each direction and a small cross-signal in the other load cell. **(d)** The slope of the signal did not depend on the location on the post that the load was placed. After calibration, residuals were small for vertical signals and slightly larger for horizontal signals for both vertical **(e)** and horizontal **(f)** loads.

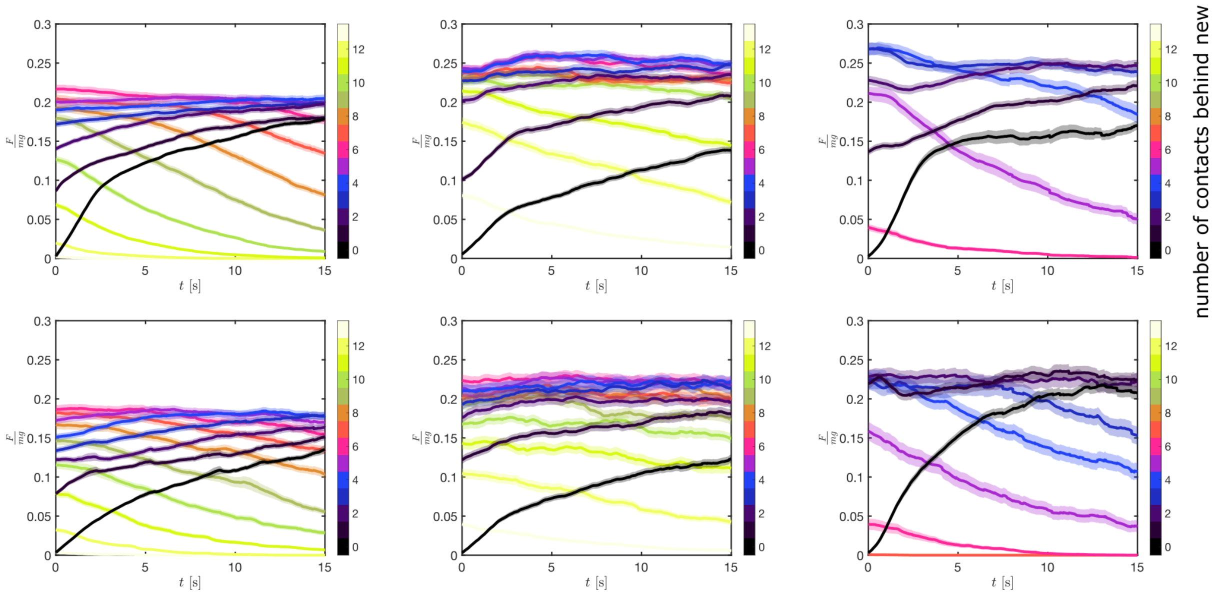


**Figure S9: Contact triggered average plots for the force magnitude on each post.** Shifting the data relative to when a new contact is formed allows us to see what the rest of the body does to accommodate this new contact.

#### S11 Contact triggered average plots

Each time the snake established contact with a new post meeting our distance and force thresholds, we marked it as an event. We then shifted force data on the new post and each of the posts contacting further down the body relative to the start of this event. The average signal on each contact could then be tracked over time. We found that across trials, after a brief period of rearrangement, the force magnitude on each post ramped up to a stable, characteristic value. This value was slightly higher and more variable with shorter posts but still stereotyped across trials. With a larger spacing there were fewer contacts to work with meaning a greater force had to be applied ot each to support the snake's weight.
